# *De novo* synthesis of a conjugative system from human gut metagenomic data for targeted delivery of Cas9 antimicrobials

**DOI:** 10.1101/2023.05.10.540216

**Authors:** Thomas A. Hamilton, Benjamin R. Joris, Arina Shrestha, Tyler S. Browne, Sébastien Rodrigue, Bogumil J. Karas, Gregory B. Gloor, David R. Edgell

## Abstract

Metagenomic sequence represents an untapped source of genetic novelty, particularly for conjugative systems that could be used for plasmid-based delivery of Cas9-derived antimicrobial agents. However, unlocking the functional potential of conjugative systems purely from metagenomic sequence requires the identification of suitable candidate systems as starting scaffolds for *de novo* DNA synthesis. Here, we developed a bioinformatics approach that searches through the metagenomic ‘trash bin’ for conjugative systems present on contigs that are typically excluded from common metagenomic analysis pipelines. Using a human metagenomic gut dataset representing 2805 taxonomically distinct units, we identified 1598 contigs containing conjugative systems with a differential distribution in human cohorts. We synthesized *de novo* an entire *Citrobacter* spp. conjugative system of 54 kb and containing at least 47 genes, pCitro, and found that pCitro conjugates from *Escherichia coli* to *Citrobacter rodentium* with a 30-fold higher frequency than to *E. coli*, and is compatible with *Citrobacter* resident plasmids. Mutations in the *traV* and *traY* conjugation components of pCitro inhibited conjugation. We showed that pCitro can be re-purposed as an antimicrobial delivery agent by programming it with the TevCas9 nuclease and *Citrobacter* -specific sgRNAs to kill *C. rodentium*. Our study reveals a trove of uncharacterized conjugative systems in metagenomic data and describes an experimental framework to animate these large genetic systems as novel target-adapted delivery vectors for Cas9-based editing of bacterial genomes.

## Introduction

Metagenomic sequencing has revealed the diversity of microbial species that compose the human microbiome and the widespread occurrence of antimicrobial resistance (AMR) genes and mobile elements that can spread AMR^1–3^. The development of multi-drug resistant bacteria has reduced the efficacy of traditional antibiotic strategies, necessitating alternative approaches to combat AMR and microbial dybioses^4, 5^. Recently, Cas9 and related nucleases derived from the clustered regularly interspaced short palindromic repeat (CRISPR) system have been repurposed as antimicrobial agents to target specific bacteria for elimination based on the introduction of double-strand breaks in the chromosome that lead to replication fork collapse and cell death^6–8^. A key unresolved issue in using Cas9 as an antimicrobial tool is delivery. Current strategies that use phage, phagemids, or conjugative plasmids as delivery vectors are based on characterized systems that replicate in *Escherichia coli* or other model bacteria and that are amenable to genetic manipulation. Delivery vectors are limited in their application by a number of factors, including host range and stability in natural (non-domesticated) bacteria and, in the case of phage vectors, development of resistance^9^. Metagenomic data represents an untapped source of potential mobile elements, and conjugative systems in particular, that could be re-purposed as host-adapted antimicrobial delivery tools.

Conjugative plasmids are well suited as delivery vectors for CRISPR nucleases or other genetic tools; they have a large coding capacity, they encode factors that promote biofilm formation to enhance cell-to-cell contact and DNA transfer^10^, they are generally resistant to restriction modification systems^11^, they do not require a cellular receptor that would provide a mechanism for bacterial resistance^12^, and they can be engineered to carry replication origins that function in a range of bacteria^13^. Conjugative systems initiate the unidirectional transfer of DNA from a donor to a recipient cell through the actions of the relaxase protein (MOB) that nicks DNA at a defined origin of transfer sequence (*oriT*)^14, 15^. Subsequent interaction of the relaxosome with the type IV coupling protein (T4CP), mating pair formation and type 4 secretion (T4SS) proteins catalyzes DNA transfer to the recipient^16–18^. Conjugative systems can exist as complete systems that are self transmissible, or as partial systems that are dependent on conjugative proteins encoded by other elements. Upon DNA transfer, some conjugative systems integrate into the chromosome as integrative conjugative elements (ICEs)^19–21^. Currently, the classification of conjugative systems is based on the identity of the MOB protein^22^.

Metagenomic sequencing coupled with bioinformatics analysis and DNA synthesis and synthetic biology optimization of gene expression can provide access to the biochemical repertoire of a community^23^. Identification of conjugative systems (and other mobile elements) from metagenomic sequencing studies where individual contigs (often generated by short-read sequencing) are assembled into metagenome-assembled genomes (MAGs) is problematic^24, 25^. Conjugative systems often have GC content that is different from the parent genome and thus would often be excluded from metagenomic binning algorithms that use GC content as a parameter for clustering. Moreover, plasmids that carry conjugative systems are not necessarily maintained in the same unit copy number as the parent chromosome causing differential sequence coverage and exclusion of the contigs that encode these systems from MAGs. Thus, our current understanding of the distribution and diversity of conjugative systems in metagenomes is likely an underestimate, and it is unknown if metagenomically-identified systems represent functional conjugation systems.

Here, we elaborate a synthetic biology approach to capitalize on the diversity of conjugative systems from human metagenomic data for delivery of CRISPR nucleases or other genetic tools. We developed a bioinformatics pipeline that searches for conjugative systems in contigs that are excluded from assembly into MAGs. Mapping of the identified conjugative systems to short-read sequences from human gut metagenomic data revealed differences in the distribution and abundances between human cohorts^1^. We selected a putative ∼54-kb conjugative system identified on a *Citrobacter* spp. plasmid and synthesized it *de novo* to create a functional conjugative plasmid, pCitro. Crucially, pCitro conjugates with a higher frequency to *Citrobacter* species than to other bacteria, is compatible with resident *Citrobacter* plasmids, and is stable over multiple generations. Programming pCitro with a TevCas9 dual nuclease and sgRNAs resulted in efficient killing of *Citrobacter*. This work provides proof of principle that conjugative systems identified in metagenomic data can be synthesized *de novo* and functionalized for the targeted killing of bacteria.

## Results

### Identification of conjugative systems from the human gut microbiome reveals differential geographical distribution

To overcome problems associated with the identification of conjugative systems from MAGs, we searched the sequences of contigs that were excluded from binning into MAGs, rationalizing that these contigs would be enriched for conjugative systems (Fig. 1A, Fig. S1). We used a dataset that represents a near complete and non-redundant picture of the human gut microbiome^1^ of 2805 taxonomically distinct units and searched 92143 contigs that did not pass quality filtering and were not binned into MAGs. Open-reading frames (ORFs) were predicted from the contigs and aligned to the UniRef90 database^26^, with the results subsequently searched by keyword for conjugation proteins. We found a total of 1598 contigs from 787 taxonomically distinct units that matched UniRef90 annotations for relaxase/MOB and T4SS/T4CP proteins (Table S1, Data S1). From these contigs, 3216 subregions where conjugative protein annotations were concentrated on the contig were extracted, retaining 2413 that were > 1kb in size (Fig. 1A). These contigs (the core conjugative contigs) represent an aggregate picture of the distribution of conjugative systems because the human gut microbiome data used here itself is an aggregate of short-read metagenomic data from many different samples.

**Figure 1.**
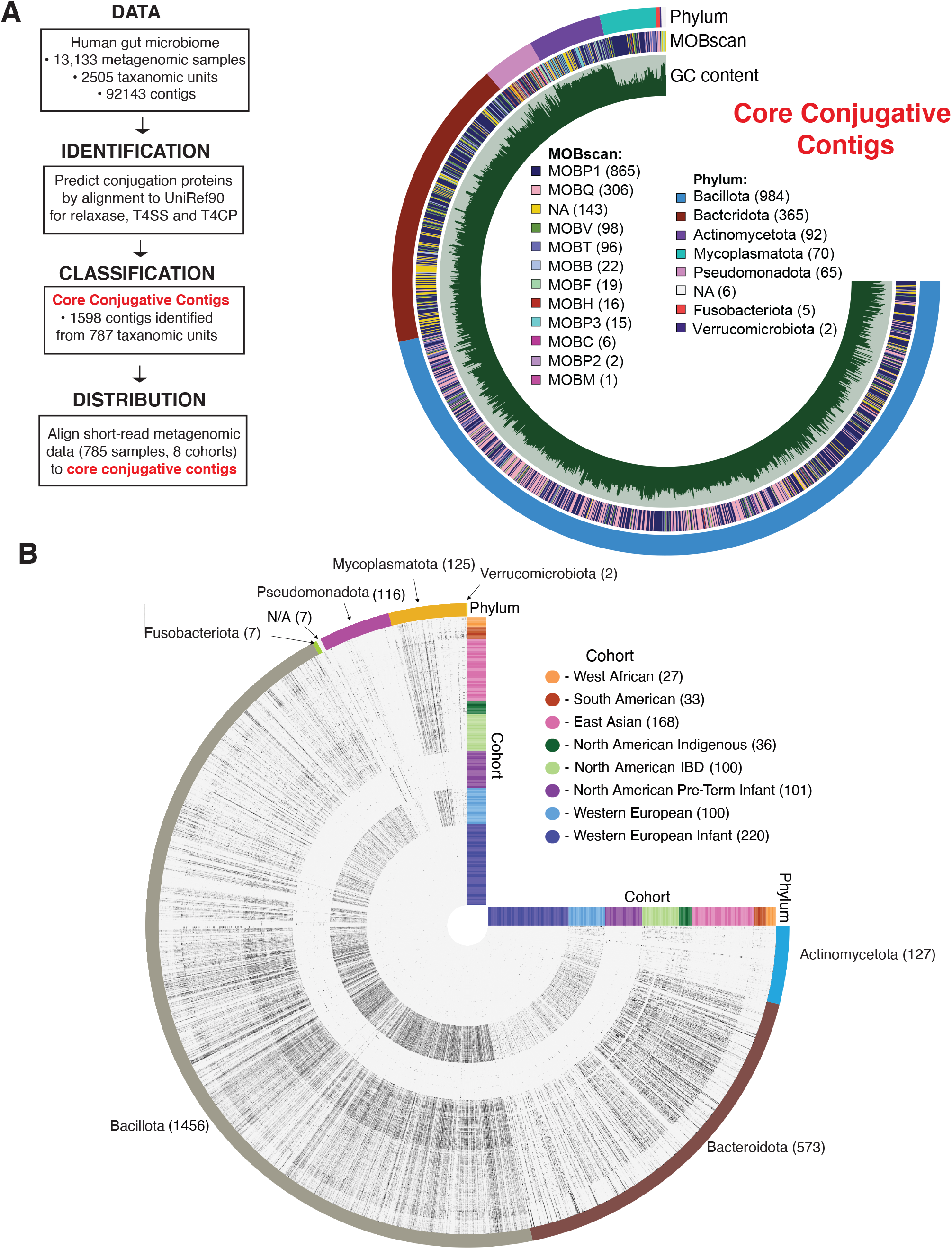
Identification and distribution of conjugative systems in human gut metagenomic data. A) Workflow (on left) for identifying conjugative systems from metagenomic data. On right is a classification of identified conjugative systems by bacterial phylum (outer ring), by relaxase (MOB) homology (middle ring), with GC-content shown on inner ring. B) Distribution of identified core conjugative contigs in different human cohorts shown by an Anvi’o plot. The outer ring denotes bacterial phylum, and the vertical and horizontal color bars indicate the human cohort. The inner rings represent the abundance of a conjugative system in each cohort as indicated by a black dot (plotted on a log2 scale), with darker points representing a higher relative abundance of a conjugative system. A file of the sequences of identified contigs is available at https://figshare.com/authors/David_Edgell/15442121

Using MobScan^27^ we predicted the mobility (MOB) families of the relaxases that were present on the identified conjugative systems (Fig. 1A, Fig. S2, Table S1). All 9 of the relaxase Mob families were present but, interestingly, MOBP1-type relaxases were found to be the majority with 865 represented systems. We used PlasFlow^28^ to predict whether the conjugative systems were encoded on a plasmid (Fig. S2, Table S1) and found that 57 were predicted to be plasmid-based. Of the remaining systems, 1353 were predicted to be chromosomal and the genetic location of 179 conjugative systems could not be determined.

To provide a higher-resolution picture of conjugative systems in different human cohorts, we aligned short-read sequencing data from 785 samples from 8 different cohorts to the core conjugative contigs (Fig. 1B, Table S2, S3). This approach allowed us to determine the abundances of each conjugative system relative to all other systems based on the read counts in the short-read sequences. Surprisingly, we found that some conjugative systems were common to all cohorts whereas other conjugative systems were private to specific cohorts. This was most notable for the North American Pre-term Infant cohort that was almost exclusively composed of conjugative systems from 16 different genera of Pseudomonadota.

In contrast, these conjugative systems were essentially absent from the Western European Infant cohort.

### Constructing a conjugative system *de novo*

The analyses in Fig. 1 revealed a significant diversity of conjugative systems in the human gut metagenome, but whether these represent functional (ie. active) conjugative systems was unknown. To address whether the putative conjugative systems were functional, we selected a 53.7 kb conjugative system (c_000000000801, hereafter called 20298) with a MOBP1 family relaxase that was extracted from a larger ∼150-kb *Citrobacter* contig for DNA synthesis (Fig. 2A, Table S1, Table S2, Data S2, Data S3). We chose this system because the ∼150-kb contig appeared to be of plasmid origin as it contained genes for a toxin/antitoxin system and a *repA* homolog). The 20298 conjugative system, or system(s) similar to support read mapping using our stringent parameters, was found in virtually every human cohort in Fig. 1C, but under-represented in indigenous populations and absent from infant populations not exposed to antibiotics (Fig. S3). This distribution suggests that the 20298 system is functional. Moreover, *Citrobacter rodentium* is a model for enterohemorrhagic *E. coli*^6, 30–32^.

**Figure 2.**
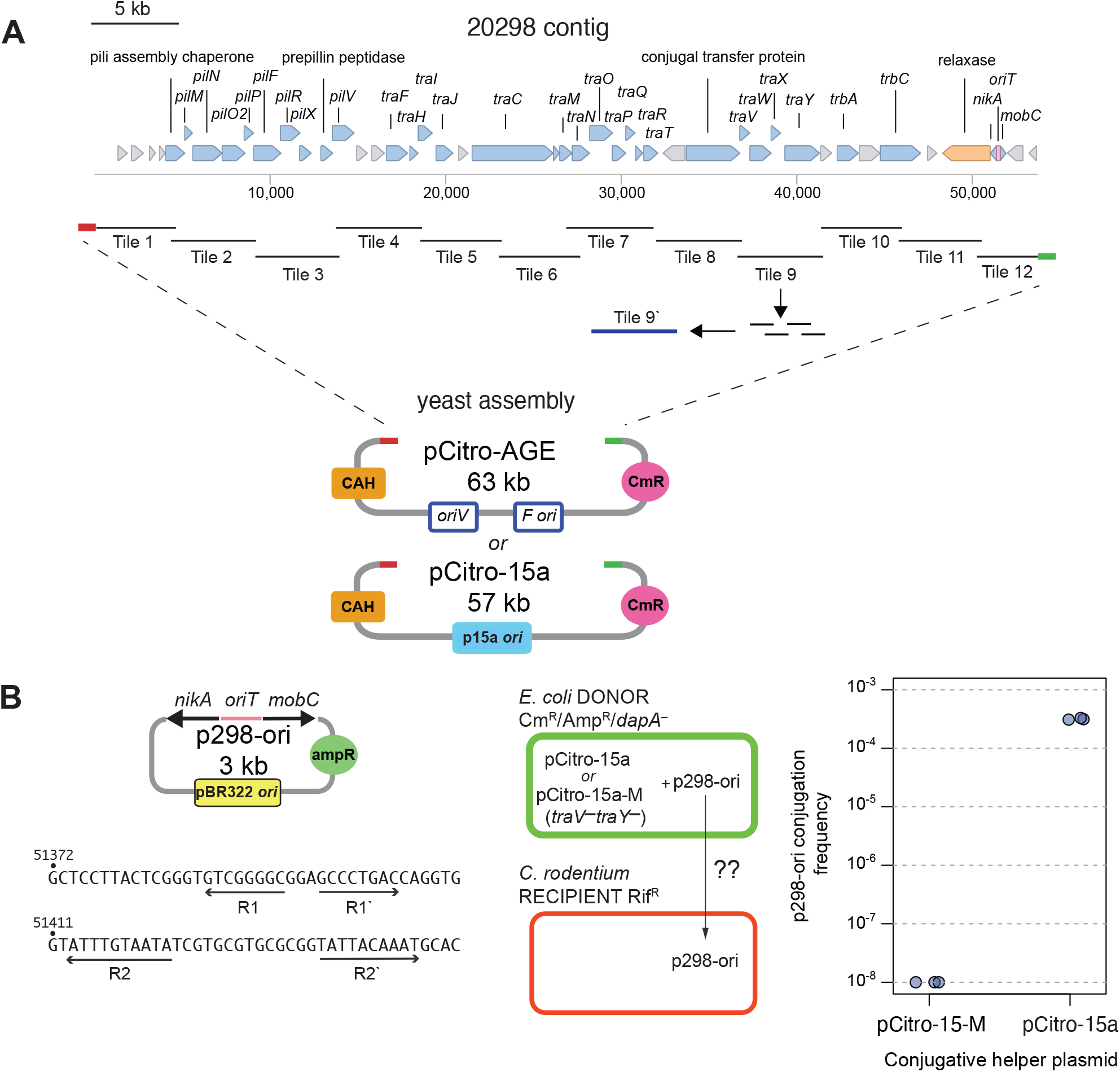
Strategy for synthesis and assembly of the 20298 contig. A) Schematic of the 20298 contig (to scale) with genes that have similarity to known conjugative functions indicated by blue-filled arrows. The putative relaxase is indicated by a orange-filled left facing arrow, and the putative *oriT* sequence by a pink rectangle. The contig was split into tiles for DNA synthesis as indicated. Tile 9 was further split into 4 smaller synthetic fragments that were assembled to create a complete tile 9’. Tiles 1 and 11 had homology hooks (red and green rectangles) for assembly into the pAGE or p15a vectors by *in vivo* yeast assembly. B) Identification and functional testing of the 20298 *oriT* sequence. The putative oriT sequence and flanking *nikA* and *mobC* genes were cloned to create p298-ori. Below the plasmid map are direct repeats identified in the 20298 *oriT* sequence^29^. At right is the strategy to test conjugation of p298-ori from *E. coli* to *C. rodentium*. pCitro-15a is a sequence verified clone of the 20298 conjugtiave contig and pCitro-15-M has inactivating mutations in the*traV* and *traY* genes. The plot shows the conjugation frequency of p298-ori from *E. coli* harbouring either pCitro-15 or pCitro-15-M to *C. rodentium*. Data points are three biological replicates.

To synthesize the 20298 conjugative system for assembly into destination plasmids by *in vivo* yeast assembly^33, 34^, we divided the 53.7-kb contig into 11 tiles of ∼5-kB that were commercially synthesized (as described in the Methods section; Data S3). The end tiles (1 and 11) contained “homology hooks” to the destination plasmids and the internal tiles had a minimum of 100-bp homology overlap with the adjacent tile (Fig. 2A). One tile (number 9) that included the coding regions for *traV, traW, traX* and *traY* could not be synthesized commercially, and was subsequently synthesized in house on a Codex DNA BioXP 3200 printer as 4 smaller fragments that were assembled into a complete tile (tile 9^′^) (Data S3). At the same time, we constructed a vector (p298-ori) that contained the putative origin of transfer (*oriT*) for the 20298 conjugative system (Fig. 2B).

We first assembled the 20298 contigs into the pAGE1^35^ backbone to create pCitro-AGE and screened for correct assembly by diagnostic restriction digest and then by whole plasmid sequencing (Fig. S4 and Table S6). Of the five clones sequenced, all contained frameshift mutations in *traV* (pilus assembly) and *traY* (Integration Host-Factor (IHF) association and *oriT* nicking) suggesting either a systematic issue with DNA synthesis of this region or that these genes were toxic in *E. coli* when this particular DNA tile was cloned independent of the adjacent tiles. Attempts to conjugate one clone (pCitro-AGE-5) from an *E. coli* donor to an *E. coli* or *C. rodentium* recipients yielded no transconjugants (Fig. 3A). We considered two reasons why conjugation of pCitro-AGE-5 was unsuccessful. First, mutations in the *traV* and *traY* genes could abolish conjugation of pCitro-AGE to both *E. coli* and *C. rodentium*. It was also possible that clones contained mutations (other than obvious frameshifts) in essential conjugation genes, but that these mutations would be difficult to identify based solely on the sequence of the metagenomically-assembled 20298 contig. Second, while the *oriV* and *F* ^′^ origins of the pAGE backbone function in *E. coli*, they may not support replication in *Citrobacter*, possibly due to incompatibility with resident plasmids. We subsequently identified 4 resident plasmids by sequencing of our *C. rodentium* strain.

**Figure 3.**
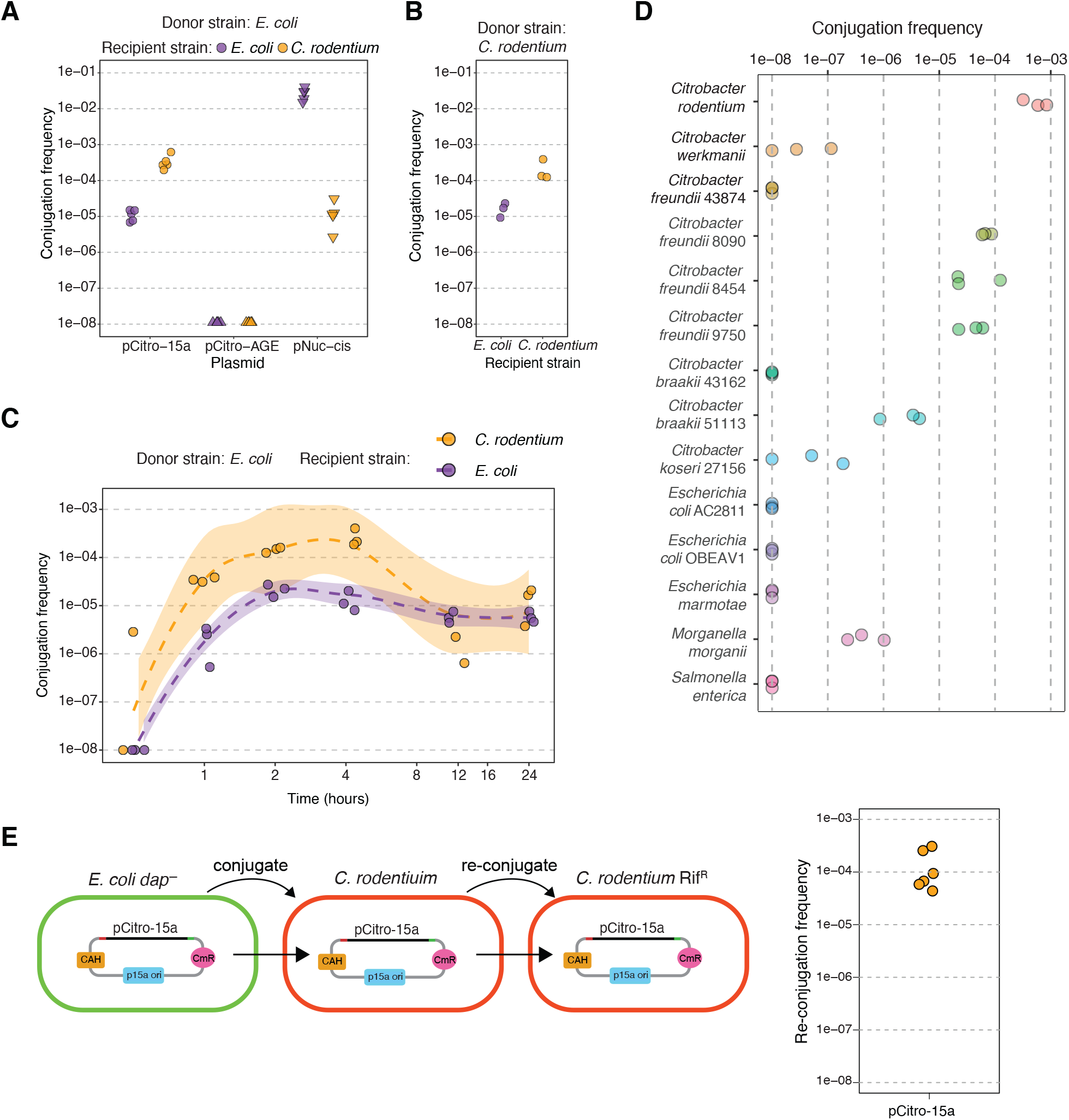
Conjugation of pCitro variants. A) Conjugation of pCitro-15a, pCitro-AGE and pNuc-cis from an *E. coli* donor to an *E. coli* or *C. rodentium* recipient. B) Conjugation of pCitro-15a from an *C. rodentium* donor to an *E. coli* or *C. rodentium* recipient. For panels (A) and (B) points represent biological replicates. C) Timecourse conjugation from 30 minutes to 24 hours of pCitro-15a from an *E. coli* EPI300 donor to an *E. coli* and a *C. rodentium* recipient. Individual dots represent biological replicates and the shaded regions indicate the 95 % confidence interval. D) Conjugation of pCitro-15a from *E. coli* to different bacteria. Each point represents a biological replicate. E) Strategy for testing sequential conjugation of pCitro-15a (on left) with frequency of conjugation from a *C. rodentium* Rif^S^ donor to a *C. rodentium* Rif^R^ recipient shown on the right. Each data point represents a biological replicate.

To resolve these issues, we attempted to complement the functions of pCitro-AGE *in trans* with different conjugative plasmids that could replicate in *E. coli* with pCitro-AGE (Fig. S5). We found one conjugative plasmid, MIP231, belonging to the IncY incompatibility group that promoted conjugation of pCitro-AGE to an *E. coli* recipient with a conjugation frequency of ∼10^−4^ – 10^−5^ (Fig. S5). Second, we used electroporation rather than conjugation to move pCitro-AGE or pAGE1 into *C. rodentium* but could not recover transformants for either plasmid, whereas transformants were readily obtainable using a p15a-based plasmid. Collectively, these experiments showed that the inability of pCitro-AGE to conjugate was due to mutations in the *traV* and *traY* genes because the pAGE1 origin did not support replication in *C. rodentium*. Using pCitro-AGE-5 as a template, we re-amplified the 20298 conjugative system using PCR primers that simultaneously corrected the *traV* and *traY* genes. These new fragments were then assembled into the p15a backbone in yeast to generate pCitro-15a (Fig. 2A and Table S7). Additionally, we constructed pCitro-15a-M that changed the plasmid backbone but maintained the inactivating mutations in the *traV* and *traY* genes that were present in pCitro-AGE.

### Preferential conjugation of pCitro-15a to *Citrobacter* strains

Using an *E. coli* EPI300 *dap-*auxotrophic donor strain we successfully conjugated pCitro-15a to *E. coli* and *C. rodentium* recipients (Fig. 3A). Crucially, conjugation from the *E. coli* EPI300 donor occurred at an approximately 30-fold higher frequency to *C. rodentium* than to *E. coli* after a 1-hour conjugation (p-value = 1.1e-06, Welch’s two-sided t-test). We obtained a similar result when we conjugated pCitro-15a from a *C. rodentium* donor to either *E. coli* EPI300 or to a rifampicin-resistant *C. rodentium* recipient; a 13-fold higher conjugation frequency to *C. rodentium* than to *E. coli* was observed (p-value = 0.008, Welch’s two-sided t-test). We also performed a timecourse conjugation experiment with the *E. coli* EPI300 donor and an *E. coli* or *C. rodentium* recipient over 24 hours (Fig. 3B). A higher frequency of conjugation to *C. rodentium* than to *E. coli* was observed over the first 4 hours, whereas the 12- and 24-hour time points showed similar conjugation frequencies. We also found that pCitro-15a had a higher frequency of conjugation from *E. coli* to *C. rodentium* than pNuc-*cis*, an IncP RK2-based conjugative plasmid that we previously constructed for Cas9-killing of *S. typhimurium*^36^. However, pNuc-*cis* had a much higher frequency of conjugation from an *E. coli* donor to an *E. coli* recipient than did pCitro-15a. We used pCitro-15a to confirm that the p298-ori plasmid (with the putatively identified origin of transfer sequence) could be conjugated *in trans* from *E. coli* to *C. rodentium*, whereas the pCitro-15-M plasmid with mutations in the *traV* and *traY* genes did not support conjugation of p298-ori (Fig. 2B).

To examine the host-range of pCitro-15a within the *Citrobacter* genus, we performed conjugations of pCitro-15a to seven additional *Citrobacter spp*. strains (Fig. 3D). After a standard one hour conjugation from the *E. coli* Epi300 donor, moderate conjugation frequencies between 2.9 × 10^−6^ and 7 × 10^−6^ were found for four of the seven strains. The remaining three strains yielded minimal or no transconjugants. We also tested conjugation to other Gammaproteobacteria, finding moderate conjugation to *Morganella morganii* but not to other bacteria. Conjugation was performed to *C. rodentium* in parallel and occurred at a higher frequency (5.9 × 10^−4^) than to any of the other species tested. We also showed that pCitro-15a could be re-conjugated from *C. rodentium* to a rifampicin-resistant *C. rodentium* recipient with a frequency similar to that observed for *E. coli* to *C. rodentium* conjugation experiments (Fig. 3E). We analyzed 4 clones each of 6 original pCitro-15a *C. rodentium* transconjugants that had been passaged on solid and liquid media by restriction digests, finding no evidence of gross rearrangements in pCitro-15a (Fig. S6). The same analyses revealed restriction fragments diagnostic of the resident *C. rodentium* plasmids. Further evidence of plasmid function and stability was obtained by mapping reads from RNA-seq of RNA extracted from *C. rodentium* or *E. coli* harbouring pCitro-15a (Fig. S7 and Table S8). In both cases, reads could be mapped to all annotated genes in the 20298 contig, but we noted a higher abundance of reads that mapped to the coding versus the non-coding strands for RNA isolated from *C. rodentium*. This differential mapping could reflect more stringent regulation by *Citrobacter* -specific regulatory elements.

Collectively, this data shows that we successfully synthesized and assembled a functional conjugation system that was identified from metagenomic data. The 20298 conjugative system functions outside of the context of the large contig on which it was identified. Crucially, the pCitro-15a plasmid has a significantly higher frequency of conjugation to *Citrobacter* species than to *E. coli*, implying that the 20298 conjugation system is optimized for the *Citrobacter* genus. Moreover, pCitro-15a appears to stably replicate in *C. rodentium* and is compatible with resident plasmids.

### Functionalizing pCitro to kill *C. rodentium* with TevCas9

To facilitate cloning of genetic cargo into pCitro-15a, we installed a destination cassette for gateway recombination cloning^37^ to generate pCitro-dest (Fig. 4A). We also created pENTR-TC9, an entry plasmid for pCitro-dest that contains the TevSpCas9 dual nuclease under the control of an arabinose-inducible promoter and a sgRNA expression cassette driven by a constitutive promoter derived from the tetracyline resistance gene (Fig. 4A). Together, these plasmids allowed us to create pCitro-dest clones with different sgRNAs targeting the *C. rodentium* genome and where correct assembly could be quickly confirmed by simple diagnostic restriction digests (Fig. 4A).

**Figure 4.**
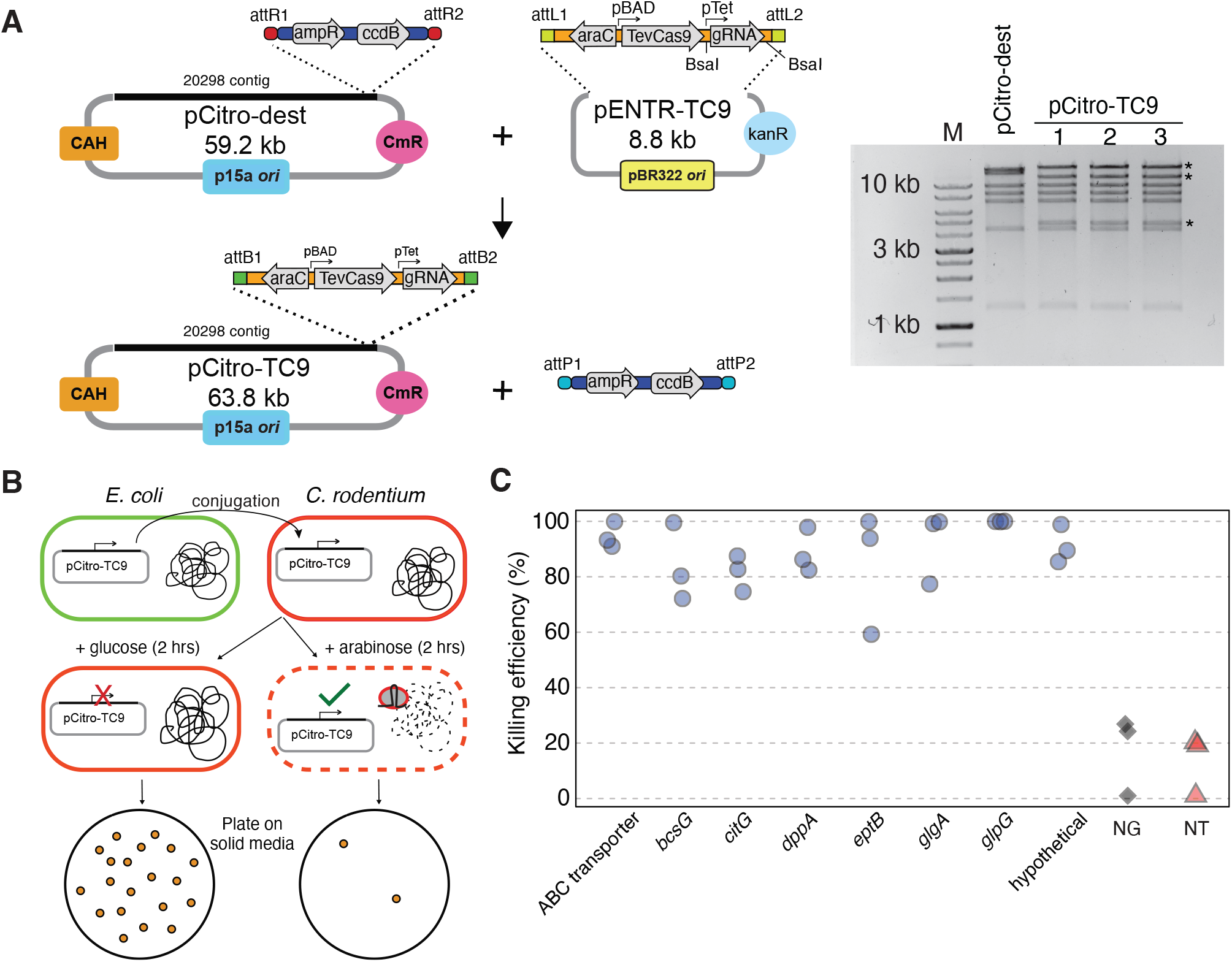
Functionalizing pCitro to kill *C. rodentium*. A) pCitro was modified to include a gateway destination cassette for cargo insertion to create pCitro-dest. pENTR-TC9 was constructed to facilitate the recombination of TevCas9 and sgRNA into the pCitro-dest plasmid. B) Mechanism of chromosomal targeting by a conjugated pCitro-TC9 plasmid. pCitro-TC9 is conjugated from *E. coli* to *C. rodentium*. TevCas9 targeting the *C. rodentium* genome is either induced with arabinose, or repressed with glucose. Transconjugants are plated and grown overnight and killing efficiency is calculated by comparing CFUs. C) Killing efficiency of *C. rodentium* with pCitro-TC9. 8 sgRNAs targeting the *C. rodentium* genome (blue circles), a non-targeted sgRNA (red triangles), and an empty sgRNA cassette control (NG, black diamonds) were tested in triplicate. Killing efficiencies were calculated as described above and expressed as a percent.

We cloned eight sgRNAs targeted to the *C. rodentium* genome that were predicted to have high activity^38^ and one sgRNA not targeted (NT) to the *C. rodentium* genome into pCitro-dest (correctly assembled clones were called pCitro-TC9) (Fig. 4B and Table S9). We conjugated pCitro-TC9 from *E. coli* to *C. rodentium* with glucose supplementation to repress TevSpCas9 expression. Individual transconjugants were grown in liquid cultures under inducing (+ 0.2% arabinose) or repressing (+ 0.2% glucose) conditions, and then plated on solid media. We determined the killing efficiency by comparing colony counts between the two conditions (Fig. 4B) and observed killing efficiences of 82-99.9% for all sgRNAs targeted to the *C. rodentium* genome (Fig. 4C). In contrast, killing efficencies for the non-targeted sgRNA, as well for pCitro-TC9 with no sgRNA (NG), ranged from 0-20% (Fig. 4C), suggesting that expression of TevSpCas9 by itself was partially toxic. Taken together, this data shows that pCitro can successfully deliver a TevSpCas9 nuclease for targeted killing of *C. rodentium*.

## Discussion

Strategies to access the functional potential of metagenomic sequences can be broadly classified into untargeted approaches that involve functional- or sequence-based screening of metagenomic DNA libraries for activities of interest^39^, or targeted approaches that identify candidate genes based on sequence or structural similarity for DNA synthesis and characterization. Indeed, a number of studies have utilized the latter, synthetic biology approach to functionalize single genes, small biosynthetic pathways, or regulatory elements from reference databases or metagenomic surveys^40–43^. Our strategy of searching contigs typically excluded from metagenomic assembly pipelines identified a trove of uncharacterized, multi-gene conjugative systems in the human gut metagenome as starting scaffolds for DNA synthesis and assembly. To our knowledge, our study is the first to reanimate a large multi-gene system *de novo* from metagenomic data.

Recent studies have mined the reference sequence databases of bacterial isolates to highlight the diversity of plasmid-based conjugation systems^44, 45^. Thus, a primary motivation for our study was to broadly reveal the diversity of uncharacterized conjugative systems in metagenomic data that could be used as scaffolds for DNA synthesis rather than to provide an exhaustive description of conjugative systems. Two notable findings arose from our analyses.The first was that conjugative systems are differentially distributed between human cohorts, which could reflect sampling bias or differences in diets, antibiotic exposure or other environmental differences that could influence bacterial composition^46, 47^. The second was the differential distribution of conjugative systems in bacterial phyla, particularly the low relative abundance of systems in Pseudomonadota relative to those found in Bacillota and Bacteroidota. This difference is striking because many previously characterized plasmid-based conjugative systems were isolated because they could replicate in, and conjugate to, model bacteria (often Pseudomonadota) cultivable in a laboratory environment. However, by using the synthetic biology strategy outlined here, uncultivable plasmid-based conjugative systems could be animated in *E. coli* (or other hosts). Interestingly, only 3.5% of identified systems were classified as plasmid-based, reflecting the limitations of current annotation tools, while 85% were classified as chromosomal that likely included integrated-conjugative elements. In the future, identification of plasmid-based conjugation systems from metagenomic communities could be enhanced by using long-read sequencing technologies, or a hybrid of long- and short-read sequencing^48^, to capture the sequences of intact plasmids^49^, as opposed to short-read sequencing that was primarily used to generate the human gut metagenome and that is not well-suited to capturing plasmids or other sequences with differential copy number^50^.

While significant, the cost of DNA synthesis for the ∼53-kb 20298 contig (∼0.11 cents per base at project start) was not a serious bottleneck. Rather, the repeated occurrence of frameshift mutations in tile 9, specifically in the *traV* and *traY* genes, was problematic. This observation suggested either a systematic issue with DNA synthesis of this tile, or that these conjugation components were toxic in *E. coli* when cloned on a multi-copy vector. Studies with the *E. coli* F plasmid have shown that TraY interacts with IHF to promote DNA bending and stimulate relaxase nicking of the *oriT* sequence^51^. TraY toxicity could be due to titration of IHF from *E. coli* regulatory sequences. TraV is involved in assembly of the conjugative pilus^52^, but potential mechanism(s) of TraV toxicity are not clear. Regardless, the use of compatible conjugative plasmids to complement inactive pCitro assemblies suggests a general strategy to ascertain whether assembled conjugative systems are functional, particularly those that do not contain obvious errors that create frameshifts or other mutations. Moreover, by using plasmid libraries of single-genes from known, functionally active systems, it would be possible to narrow down inactivating mutations on the assembled plasmids. Alternatively, the use of a different ‘storage host’ for the synthesized tiles, such as *Saccharomyces cerevisiae* or *Bacillus subtilis*^53, 54^, could avoid issues of toxicity.

Our data imply that target-adapted conjugative systems are abundant in the human gut metagenome. Interestingly, we observed a higher frequency of conjugation of pCitro-15a to *C. rodentium* than to other bacteria, agreeing with other studies that showed specificity in the conjugation process^55, 56^. A number of past studies have utilized conjugative systems with broad host ranges in an attempt to access a significant fraction of the targeted microbiome^57, 58^. While promising, low conjugation rates to diverse bacteria in the targeted microbiome and stability of conjugative plasmids with resident plasmids are confounding factors. In many cases, conjugative systems required optimization by *in vitro* evolution^6^, gene minimization^59^, or other manipulations to enhance conjugation to desired recipients^60^. In contrast, by tapping the natural diversity of conjugative systems optimized for specific bacterial genera, or specific species, issues such as conjugation efficiency, compatibility with resident plasmids, and stability have been evolutionary pre-optimized for the microbial communities of interest.

In summary, we constructed a large synthetic genetic system *de novo* from metagenomic data that contains at least 47 predicted genes, and showed the conjugative function of the system. Codon optimization, removal of de-stabilizing repetitive sequences and toxic genes, and addition of biocontainment systems could be included in the synthetic biology strategy to fine-tune each conjugative system for its intended use. Similar approaches to those described here could be used to identify and functionalize other mobile genetic systems for microbiome editing.

## Materials and Methods

### Bacterial and yeast strains

All yeast assemblies were performed using *Saccharomyces cerevisiae* VL6-48 (*MAT* **a**, *his3*Δ*200, trp*Δ*1, ura3-52, ade2-101, lys2, psi+cir*°). *E. coli* EPI300 (F’ *λ* ^-^ *mcrA* Δ(*mrr-hsdRMS-mcrBC*) *ϕ* 80d*lacZ* Δ*M15* Δ*(lac)X74 recA1 endA1 araD139* Δ(*ara, leu)7697 galU galK rpsL* (Str^R^) *nupG trfA dhfr*) (Epicentre) as well as a diaminopimelic acid (DAP) auxotroph of the strain^35^ were used for cloning and as conjugative donors. *Citrobacter rodentium* DBS100, *Salmonella enterica* Typhimurium LT2, and kanamycin-resistant *E. coli* strain were used as primary recipients. Seven additional *Citrobacter* spp. strains were used as recipients. A complete list of plasmids and strains used can be found in Table S10.

### Synthesis and cloning of the 20298 contig

The conjugative system identified from the conjugative contig “20298” was synthesized in 11 fragments from Twist Bioscience as clonal genes (Data S3). One fragment of the sequence was additionally synthesized on a Codex BioXP 3200. Fragments ordered from Twist Bioscience were released from their vectors using a PmeI digest, while the tile constructed using the BioXP 3200 was PCR amplified. Each tile contained 86-476 base pairs of homology with the adjacent fragment. The terminal DNA fragments contained homology with the backbone of plasmid pAGE1.0^35^, which was originally derived from p0521s^61^. pCitro-AGE was constructed with these fragments using a modified yeast assembly^33, 34^, which we have previously described in detail^7^. Once it was identified that pCitro-AGE contained mutations in the *traV* and *traY* genes, and that the pAGE replicon was non-functional in *C. rodentium*, the conjugative contig was re-cloned into the backbone derived from pNuc-trans^7^. This contains a p15a origin, chloramphenicol-acetyltransferase and CEN6-ARS4-HIS3. The conjugative contig was reamplified from pCitro-AGE with PCR, using primers the fix the mutations in the *traV* and *traY* genes. The remainder of the plasmid was amplified in fragments with overlapping homology to the adjacent fragment. These fragments were then assembled in yeast assembly as described above to generate pCitro-p15a. Sequences all assembled plasmids are available in the Supplementary Data (Data S4-S9).

### sgRNA cloning

A gateway cloning donor vector was constructed into the pDONR221 backbone, containing an arabinose-inducible TevCas9 endonuclease^62^ and an sgRNA cloning cassette flanked by *attL* recombination sites to form pENTR-TC9. A corresponding gateway destination cassette containing a *ccdB* toxin gene and ampicillin resistance gene flanked by attR sites was cloned into pCitro-15a, to generate pCitro-15a-dest. We used crisprHAL^38^ to identify sgRNAs predicted to have high killing efficiency against *Citrobacter* (Table S9). We cloned eight sgRNAs (Table S5) into pENTR-TC9 using Golden Gate assembly at the BsaI cassette. A gateway LR reaction was performed and the resulting plasmid was transformed into *E. coli* EPI300 (Dap-) to insert pENTR-TC9 variants containing sgRNAs into pCitro-15a-dest, forming pCitro-TC9.

### Bacterial conjugation assays

Donor and recipient strains were grown to saturation overnight in selective media. Saturated cultures were then diluted 1:50 into 5 mL non-selective LSLB (10 g/L tryptone, 5 g/L yeast extract, 5 g/l NaCl) supplemented with 60 *μ*g/mL diaminopimelic acid (DAP) and grown to an A600 of 0.5-1.0 and adjusted to an A600 of 0.5. Cultures were centrifuged at 4000 RCF for 5 minutes and resuspended in 500 *μ*L of 10% glycerol. Cells were aliquoted and frozen at -80 ^°^C. For conjugations experiments, cells were thawed on ice and 50 *μ*L of donor strain were mixed with 50 *μ*L recipient strain on LSLB plates supplemented with 60 *μ*g/mL DAP. Conjugations proceeded on plates at 37 ^°^C for 1 hour. Cells were scraped into 400 *μ*L LSLB with a glass spreader. These cell suspensions were then serially diluted and plated on selection for donors, recipients, and transconjugants. Plates were incubated for 16-20 hours at 37 ^°^C and colonies were counted manually.

### Plasmid stability assays

pCitro-15a was conjugated to *C. rodentium* as described above. Transconjugants were isolated and passaged on selective plates with overnight growth at 37 ^°^C for 16-20 hours. Colonies were isolated from these plates and passaged once again on selective plates at 37 ^°^C for 16-20 hours. Individual colonies were picked from these plates and grown overnight in selective liquid LSLB at 37 ^°^C. Plasmids were extracted and diagnostic digests were performed with NotI-HF and NcoI-HF (New England Biolabs). Digests were analyzed on agarose gels for plasmid rearrangements.

### *C. rodentium* killing assays

pCitro-15-TC9 plasmids with sgRNAs cloned were conjugated from *E. coli* (Dap-) to *C. rodentium* on LSLB + Dap 60 *μ*g/mL + 0.2 % glucose plates at a 4:1 donor:recipient ratio for 1 hour. Cells were scraped into 500 *μ*L LSLB, and added to an additional 500 *μ*L LSLB. 250 *μ*L of the resulting cell suspension was diluted 1:2 into 2x selective and inducing LSLB (50 *μ*g/mL chloramphenicol and 0.4 % arabinose) and selective and 2x repressing LSLB (50 *μ*g/mL chloramphenicol and 0.4 % glucose). Cell suspensions are therefore at final concentrations of 25 *μ*g/mL chloramphenicol and 0.2 % arabinose for induced, and 25 *μ*g/mL chloramphenicol and 0.2 % glucose for repressed. Cells were then incubated at 37 ^°^C with shaking for 2 hours to induce TevCas9 expression. After 2 hours of growth, the suspensions were centrifuged and resuspended in LSLB supplemented with 25 *μ*g/mL chloramphenicol. Cell suspensions were then serially diluted and plated on LSLB supplemented with 25 *μ*g/mL chloramphenicol and 0.2 % glucose. Colonies were counted and killing efficiency was calculated using the following formula.

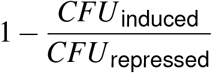

### Identifying conjugative plasmids in metagenomic datasets

A detailed workflow is available at https://github.com/bjoris33/ce_synthesis_paper. Briefly, a near-complete and non-redundant set of human gut microbiome genomes were downloaded from the European Bioinformatics Institute FTP site^1^. A set of genomes that represent a near-complete array of bacterial species capable of colonizing the human intestinal tract was downloaded from the European Bioinformatics Institute FTP site^1^. Open reading frames were predicted in the genome by Prodigal version 2.6.3^63^ and the predicted protein sequences were then aligned to the UniRef90 database^26^ using the Diamond protein aligner version 0.9.14^64^. Contigs were extracted from the genomes if they contained annotations for a relaxase/mobilization protein and a type IV secretion/type IV coupling protein using a word-search strategy. A FASTA formatted file of the 1538 conjugative contigs is available at https://figshare.com/authors/David_Edgell/15442121. Short-read data from 785 samples (Supplemental Table x)^65–72^ were downloaded from the Sequence Read Archive using the SRA toolkit version 2.9.2. Reads were deduplicated with ‘dedupe.sh’^73^, and trimmed with Trimmomatic version 0.36^74^ with options ‘LEADING:10 TRAILING:10’. Subregions of the contigs where annotations for conjugative proteins were present, with no more than 20 ORFs between successive UniRef90 annotations for conjugative proteins, were extracted. The processed read data were mapped to the extracted conjugative systems using Bowtie2 version 2.3.5^75^ with the settings ‘–no-unal –no-mixed –no-discordant’. Extraction of the sub-regions was to avoid an artificially high proportion of reads mapping in samples where the bacterial species is present, but the ICE has not integrated in its genome (Fig. S1). The proportion of reads mapping to the conjugative systems was extracted from the Bowtie2 output, and the mapping data was visualized using Anvi’o^76^. Identity as either a plasmid or chromosomal sequence was predicted using PlasFlow^28^. Relaxase mobility grouping of the cojugative element was predicted using MOBscan^27^.

### RNA-seq

Total RNA was prepared from three replicates of pCitro-15a electroporated into *E. coli* EPI300 (*dap*-) and *C. rodentium*. A single colony from each transformation was grown overnight to saturation under selection. Saturated cultures were diluted 1:50 into non-selective LSLB, and LSLB supplemented with 60 *μ*g/mL DAP for *C. rodentium* and EPI300 respectively. Cultures were grown for 2 hours at 37 ^°^C with 225 RPM shaking, reaching OD600 between 0.4-0.6. 1.5 mL of culture were centrifuged was centrifuged at 16000 RCF for 5 minutes at 4 ^°^C. The supernatant was aspirated and the pellet was resuspended in 1 mL Trizol reagent (Invitrogen). Samples were incubated at 65 ^°^C for 10 minutues, and 0.2 mL of chloroform was added and mixed by inverting for 15 seconds. Samples were incubated at room temperature for 3 minutes before being centrifuged at 16000 RCF for 5 minutes at 4 ^°^C. The aqueous phase was transferred to a clean tube and 1 volume of ethanol was added to each. An NEB Monarch cleanup kit was used by adding samples to an RNA cleanup column and centrifuging at 11000 RCF for 1 minute at room temperature. The column was washed twice by adding 500 *μ*L of RNA cleanup wash buffer and centrifuging at 11000 RCF. The column was spun for an additional 1 minute at 11000 RCF to dry before eluted into 50 *μ*L nuclease-free water by centrifuging at 11000 RCF for 1 minute.

Twelve libraries were prepared and sequenced using an Illumina NextSeq High Output 75 cycle sequencing kit on an Illumina NextSeq. Reads were trimmed with Trimmomatic version 0.36^77^ with options LEADING:10 TRAILING:10. Processed reads for *E. coli* EPI300 samples were mapped to the *E. coli* strain K-12 NEB 5-alpha genome (CP017100.1) and pCitro-15a plasmid reference sequences using Hisat2 version 2.2.0^78^. Processed reads for C. rodentium samples were mapped to the Citrobacter rodentium strain DBS100 genome (CP038008.1) and pCitro-15a plasmid reference sequences using the same workflow. Positional read coverage was determined using Samtools version 1.10^79^.

## Acknowledgements (not compulsory)

This work was supported by a Project Grant (PJT-159708) from the Canadian Institutes of Health Research to D.R.E, G.B.G. and B.J.K. T.A.H. was supported by a Postgraduate Doctoral Fellowship from the Natural Sciences and Engineering Research Council of Canada.

## Author contributions statement

T.A.H., B.R.J., B.J.K., G.B.G and D.R.E. conceived of the experiments, analyzed data and wrote the paper. A.S. performed experiments and analyzed data. T.S.B. performed bioinformatics analyses of RNA-seq data. E.A-V. and S.R. provided bacterial strains and plasmids, and wrote the paper. All authors reviewed the manuscript.

## Supplementary Information

Fig. S1 - Strategy for mapping coverage of integrated conjugative elements

Fig. S2 - Distribution of MOB family relaxases in the human gut metagenome

Fig. S3 - Abundance and distribution of the 20298 conjugative system in human cohorts

Fig. S4 - Diagnostic screening of pCitro assemblies

Fig. S5 - Complementation of pCitro with compatible conjugative plasmids

Fig. S6 - Stability of pCitro after passaging

Fig. S7 - RNAseq analysis of pCitro in *E. coli* and *C. rodentium*

Data S1 - FNA file of identified conjugative systems (available at https://figshare.com/authors/David_Edgell/15442121)

Data S2 - Sequence of the 20298 contig (GenBank format)

Data S3 - Sequences of the 20298 tiles for DNA synthesis (FASTA)

Data S4 - pCitro-AGE sequence (GenBank format)

Data S5 - pCitro-15a sequence (GenBank format)

Data S6 - pOri-298 sequence (GenBank format)

Data S7 - pCitro-TC9 sequence (GenBank format)

Data S8 - pCitro-DEST sequence (GenBank format)

Data S9 - pENTR-TC9 sequence (GenBank format)

Table S1 - Identified conjugative systems

Table S2 - Read coverage of conjugative systems in short-read data of human cohorts

Table S3 - Key for mapping short-read sample data to cohort information

Table S4 - Predicted functions of 20298 conjuative system

Table S5 - Oligonucleotides used in this study

Table S6 - Error report for pCitro-AGE assemblies

Table S7 - Error report for pCitro-15a assemblies

Table S8 - RNAseq read coverage

Table S9 - Predicted sgRNA activity scores

Table S10 - Strains and Plasmids used in this study

