## Supplementary Figures for "*De novo* synthesis of a conjugative system from human gut metagenomic data for targeted delivery of Cas9 antimicrobials"

Fig. S1 - Strategy for mapping coverage of integrated conjugative elements
Fig. S2 - Distribution of MOB family relaxases in the human gut metagenome
Fig. S3 - Abundance and distribution of the 20298 conjugative system in human cohorts
Fig. S4 - Diagnostic screening of pCitro assemblies
Fig. S5 - Complementation of pCitro with compatible conjugative plasmids
Fig. S6 - Stability of pCitro after passaging
Fig. S7 - RNAseq analysis of pCitro in *E. coli* and *C. rodentium*

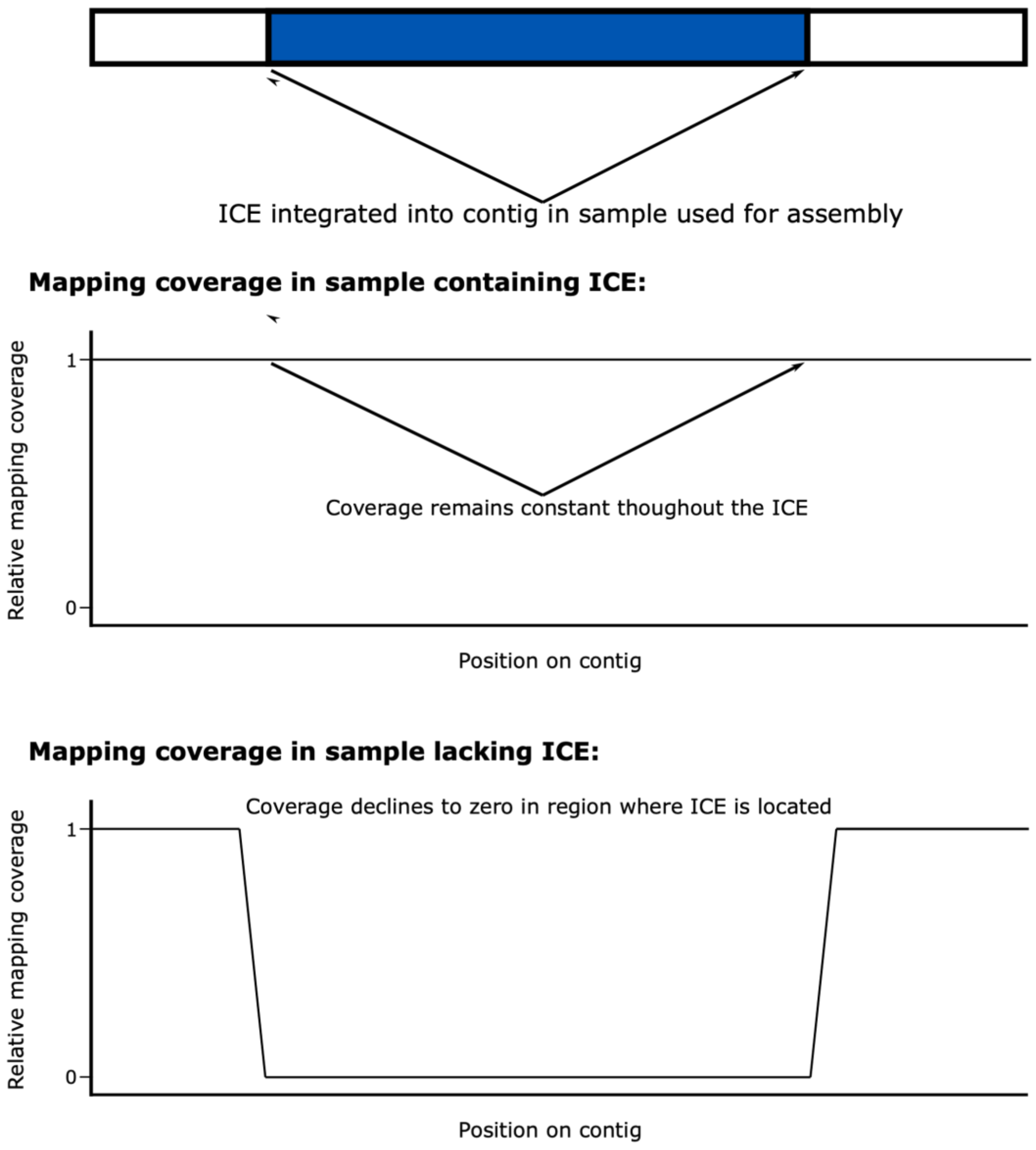


**Figure S1.** Conceptual diagram of the mapping coverage of an assembled integrative and conjugative element. The mapping coverage in the first plot shows an even mapping coverage across the contig because the ICE is present in the sample and the average mapping coverage of the contig would be an accurate metric. In the second plot, the ICE is missing in the sample and the mapping coverage falls to zero where the ICE is located on the contig and the average mapping coverage for the entire contig would be artificially high. Limiting the mapping to only the region containing the conjugative proteins solves this issue.

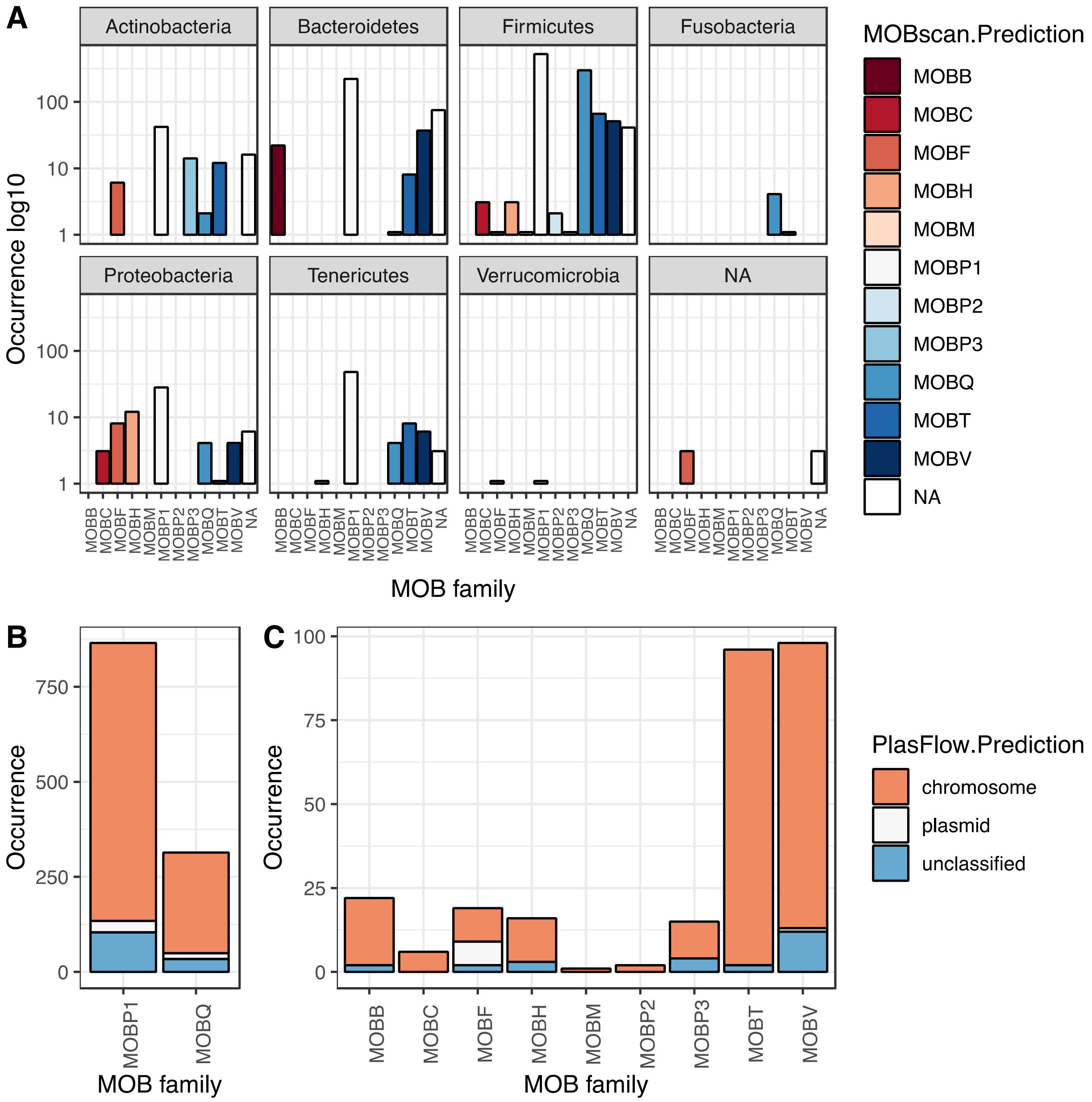


**Figure S2.** Distribution of MOB families for conjugative relaxases identified in a gut metagenomic data set. **A.** Occurrence of 11 MOB families in different bacterial phyla as predicted by MOBScan. **B.** Origin of conjugative systems as either chromosomal (orange), plasmid (white), or unclassified (blue) for MOBP1 and MOBQ families as predicted by PlasFlow. **C.** Origin of conjugative systems as either chromosomal (orange), plasmid (white), or unclassified (blue) for remaining MOB families as predicted by PlasFlow.

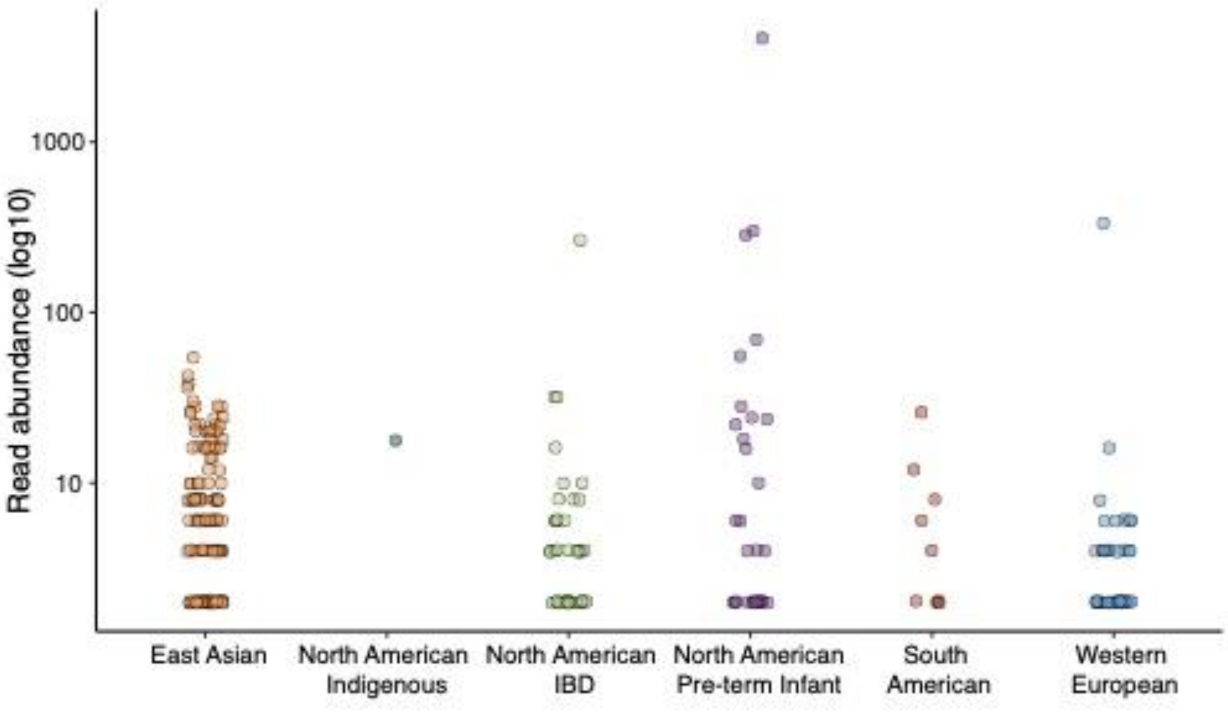


**Figure S3.** Distribution and abundance of the 20298 conjugative contig in different human cohorts. Each point represents the log10 read coverage of the 20298 contig in samples from short-read data shown in Fig. 1B.

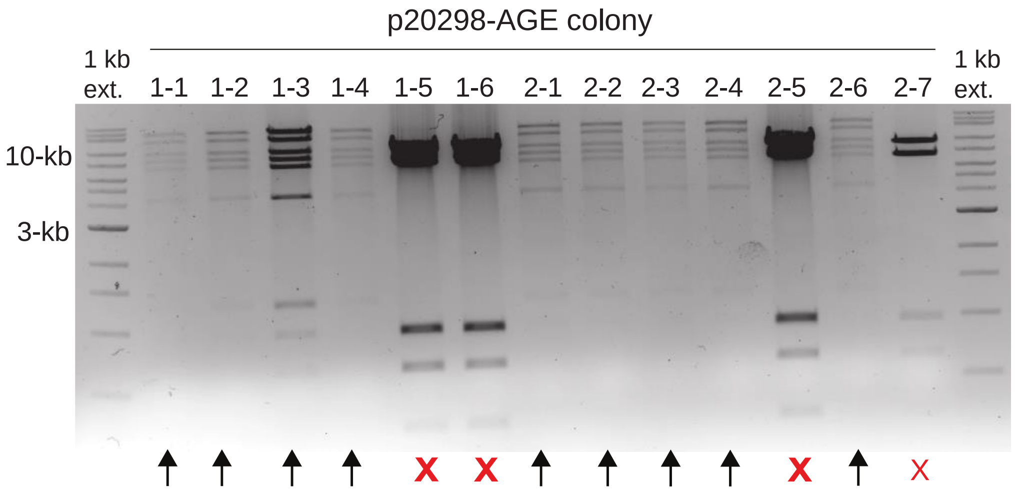


**Figure S3.** Diagnostic screening of pCitro-AGE yeast assemblies clones after transformation into *E.* coli. Isolated plasmids were digested with NotI and NcoI. Arrows indicate lanes with expected banding patterns for correct assemblies.

**A**

| Name | Inc group | Resistance | Isolated from | Size (bp) | GenBank Accession | Institute Pasteur ID |
|---|---|---|---|---|---|---|
| R64 | IncI1 | Sm, Tc | *Salmonella typhimurium* | 120,826 | NC_005014.1 | CRBIP19.30 |
| pIP112 | IncI1 | Km, Cib | *Salmonella enterica* | 100,500 | None | CRBIP19.38 |
| R69-2 | IncM | Amp, Km | *Salmonella enterica* | 78,999 | KM406488.1 | CRBIP19.53 |
| pIP69 | IncM | Ap, Km, Tc, Hg | *Salmonella enterica* | 70,500 | None | CIP pIP69 |
| pIP113 | IncN | Tc | *Salmonella enterica* | Unknown | None | CRBIP19.34 |
| pIP72 | Inc10 | Km | *Escherichia coli* | Unknown | None | CRBIP19.42 |
| pRts1 | IncT | Km, Sp | *Proteus vulgaris* | Unknown | None | CRBIP19.43 |
| pIP175 | IncI2 | Ap | *Salmonella enterica* | Unknown | None | CRBIP19.46 |
| pIP55 | IncA/C | Ap,Km,Hg,Su,Tm,Gm | *Klebsiella penumoniae* | Unknown | None | CRBIP19.51 |
| pIP16a | IncC | Ap,Km,Su | *Providencia stuartti* | Unknown | None | CIP pIP16a |
| MIP231 | IncY | Tc, H2S | *Escherichia coli* | 60,000 | None | CIP MIP231 |

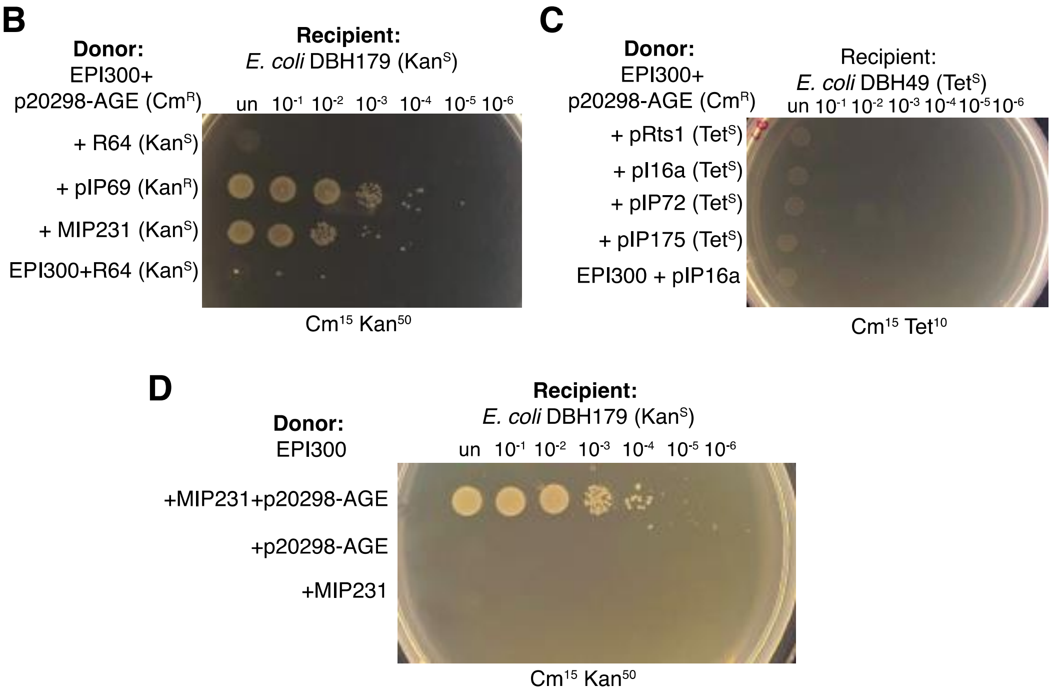


**Figure S4.** Complementation of pCitro-AGE conjugation with alternative conjugative systems**.** **A.** Information regarding tested conjugated systems for rescue. **B.** Complementation of pCitro-AGE with R64 was unsuccessful. MIP231 appears to complement conjugation. pIP69 is kanamycin resistant, obstructing the ability to determine if conjugation has occurred. **C.** Complementation of pCitro-AGE with pRts1, pI16a, pIP72, and pIP175 were unsuccessful. **D.** Validation of p20298-AGE complementation with MIP231.

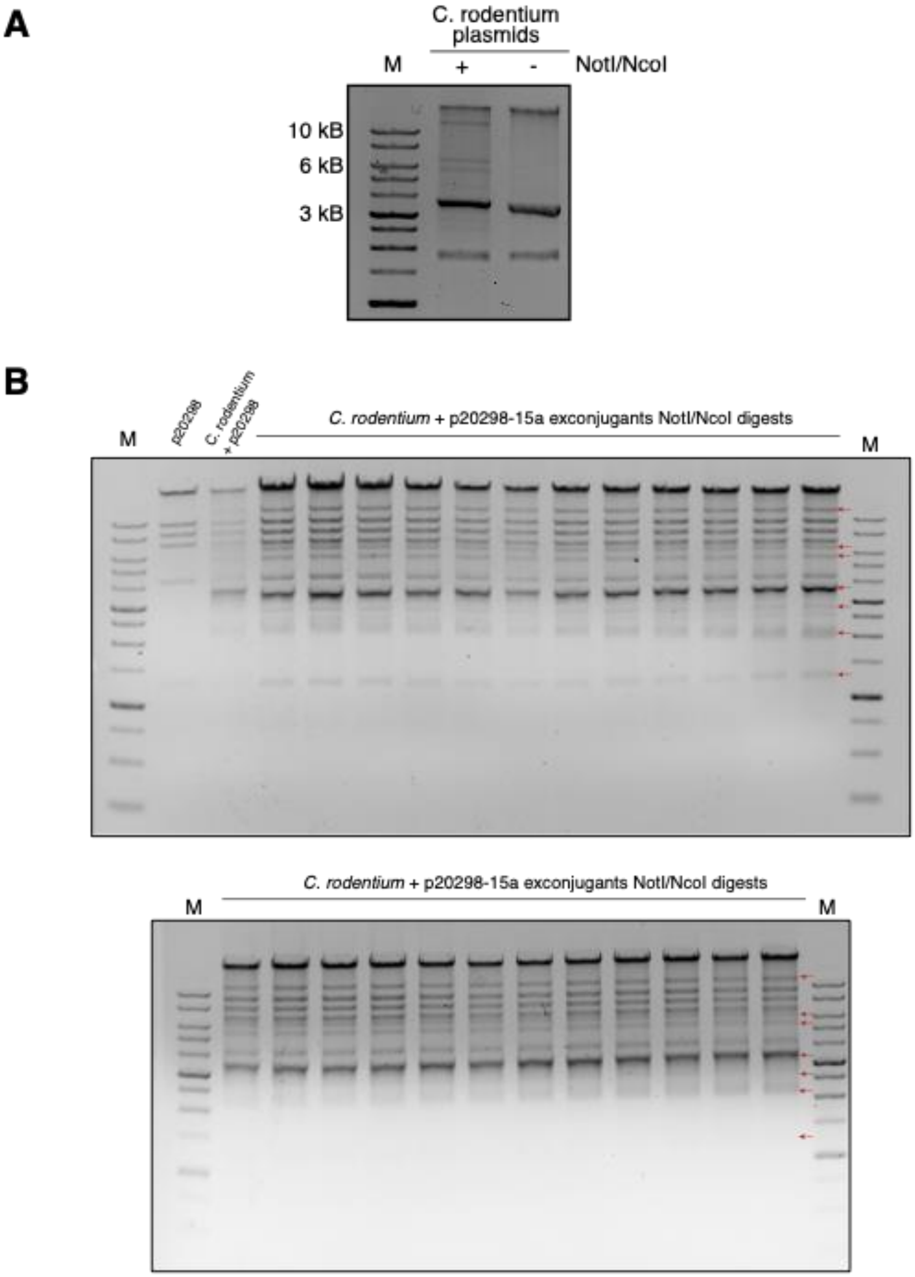


**Figure S5. A.** Diagnostic digests of native *C. rodentium* plasmids. Plasmids were extracted and digested with NotI and NcoI. **B.** Diagnostic digests of pCitro-15a transconjugants isolated from *C. rodentium* after being passaged. Colonies were passaged twice on plates before being diluted and grown to saturation overnight for plasmid extraction. Extracted plasmids were digested with NotI and NcoI.

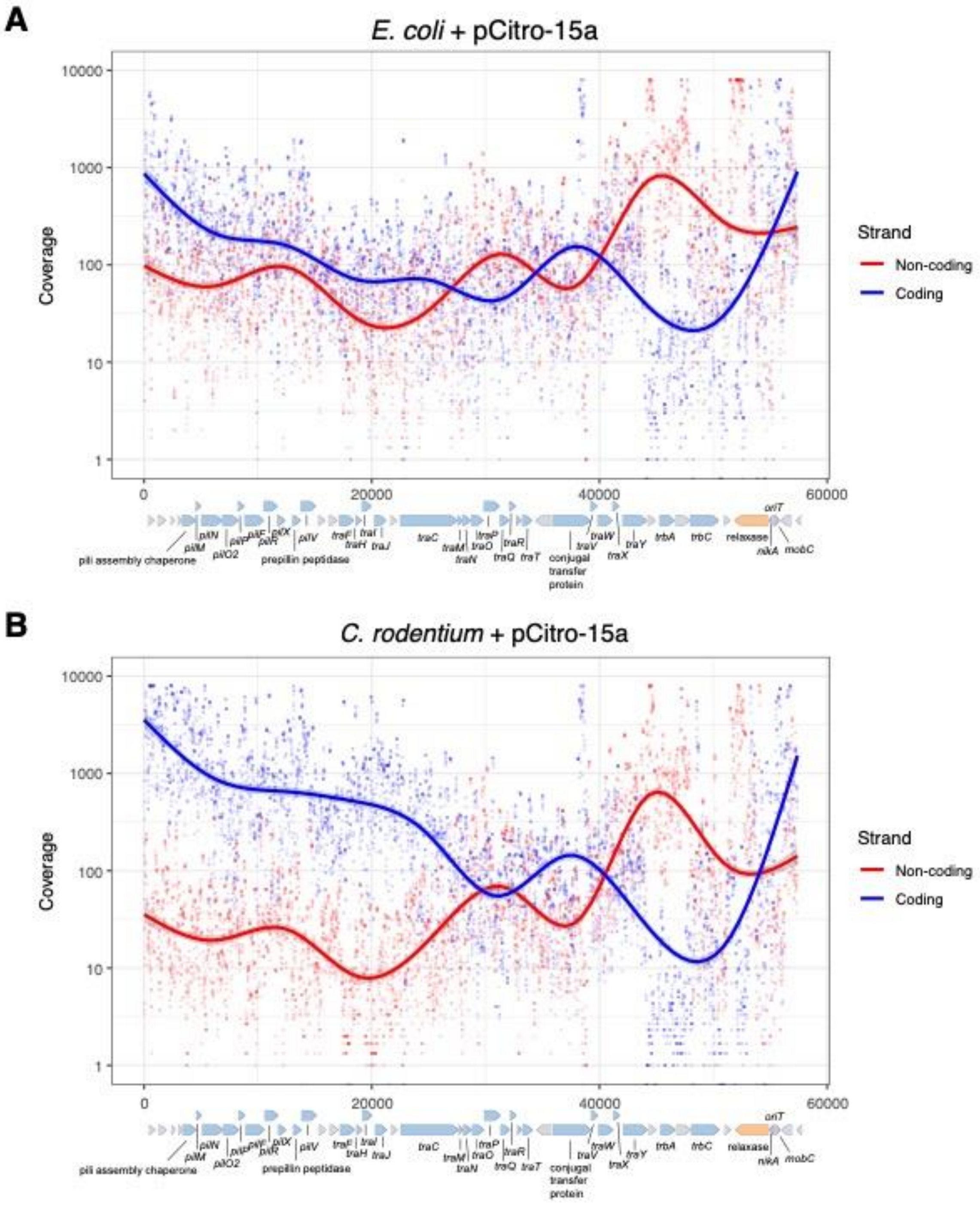


**Figure S6.** Nucleotide coverage from RNA-seq analysis of pCitro-15a expression in **A.** *E. coli* Epi300 (DAP-) and **B.** *C. rodentium*. Data is coloured based on expression from either the coding strand (blue) or non-coding strand (red).
